# Modulatory Effects of IFN-γ and IL-22 on Inflammatory Signaling and Cellular Responses in Intestinal Epithelial Cells

**DOI:** 10.1101/2024.05.09.593373

**Authors:** Asia Johnson, Timothy L. Denning

## Abstract

Inflammatory bowel disease (IBD) continues to affect millions worldwide, with an increasing prevalence that highlights the urgent need for deeper understanding of its underlying immune mechanisms. The cytokine interactions, especially those mediated by cells from the TH1 and TH17 lymphocyte subsets, are crucial in orchestrating the immune landscape of IBD. TH1 cells are well known for producing TNF-α and IFNγ, which have been extensively studied for their roles in conjunction with each other within the context of IBD. TH17 cells secrete IL-22 and IL-17, with existing studies primarily focusing on IL-22’s interaction with IL-17 rather than its interplay with other cytokines such as IFNγ. Our study focuses on the co-stimulatory effects of IL-22 and IFNγ using organoids derived from mouse small intestines to model epithelial interactions. We found that IFNγ interferes with the capacity of IL-22 to up-regulate antimicrobial peptides, which is essential in mucosal defense. Additionally, higher concentrations of IL-22 enhance IFNγ’s ability to stimulate TNF-α gene expression and CXCL10 protein production, indicating a dose-dependent relationship. This co-stimulation also led to an increased rate of cell death, influenced partly by TNF-α. These findings suggest that IL-22, typically seen as an anti-inflammatory agent, can assume a pro-inflammatory role when combined with IFNγ, complicating its effects on epithelial cells. This study highlights the need to consider specific cytokine interactions in developing more effective IBD treatments.

## Introduction

Inflammatory Bowel Disease (IBD) includes Crohn’s disease (CD) and ulcerative colitis (UC), both of which are marked by persistent inflammation within the gastrointestinal (GI) tract (McDowell, 2023). CD manifests anywhere along the GI tract but predominantly targets the small intestine, whereas UC specifically affects the large intestine. The etiology of IBD has been attributed to a complex interplay of factors, including microbiota dysbiosis, genetic predispositions, immune system dysregulation, and environmental factors (McDowell, 2023; Hu, 2022). The prevalence of IBD is notably higher in Western societies, with newly industrialized regions experiencing a rise in incidence rates (Ng et al., 2017). Lifestyle differences, such as diet, are suspected to drive this trend (Chiba, 2019).

Understanding IBD necessitates a comprehensive grasp of the immune system’s role, which is characterized by an intricate network of cytokines and chemokines. The cytokine interactions, especially those mediated by cells from the TH1 and TH17 lymphocyte subsets, are crucial in orchestrating the immune landscape of IBD. TH1 cells are well-known for producing TNF-α and IFNγ, which have been extensively studied for their roles in conjunction with each other within the context of IBD (Fish, 1999). TH17 cells secrete IL-22 and IL-17, with existing studies primarily focusing on IL-22’s interaction with IL-17 rather than its interplay with other cytokines such as IFNγ.

In healthy individuals, IL-22 is typically absent from the large intestine but is constitutively present in the small intestine (Mizoguchi, 2018). Researchers have observed elevated levels of IL-22 in samples from both UC and CD patients, where CD IL-22 related pathology is largely part of a TH1/TH17 response. In UC, the dominant relevant response is a TH2 response, and there are lower levels of IL-22 compared to a TH1/TH17 response (Mizoguchi, 2018; Abraham, 2009). IL-22 is crucial in maintaining the integrity of the epithelial barrier by stimulating mucin and antimicrobial peptide (AMP) production. This cytokine not only induces mucin production but also promotes the proliferation of goblet cells (Kier, 2020). IL-22 is vital for tissue repair following acute injury. The absence of IL-22 impairs tissue regeneration (Aparicio-Domingo, 2015), and IL-22-deficient mice exhibit increased susceptibility to Citrobacter rodentium infection, highlighting its protective role in host defense (Zheng, 2008).

IFNγ is the only type 2 interferon and is produced by TH1, CD8+ T cells, NK cells, and ILC1 (Murray, 2002; Conlon, 2021). Binding of IFNγ to its receptor activates the JAK1 and JAK2 pathways, resulting in the phosphorylation and dimerization of STAT1. These homodimers then translocate to the nucleus to bind to IFNγ activated site (GAS) elements on IFN-stimulated genes (ISGs), initiating transcription (Platanias, 2005). IFNγ is crucial for defending against intracellular bacterial and viral pathogens. Deficiencies in IFNγ significantly increase susceptibility to infections by organisms such as Salmonella and HSV-1 (Perez-Toledo, 2020; Kim, 2022).

IFNγ often plays a detrimental role in IBD. The polymorphism IFNG rs1861494 is associated with increased IFNγ secretion and more severe IBD symptoms (Gonsky, 2014). Transcriptomic studies have found elevated IFNγ gene expression in UC patients (Gao, 2022). The influence of IFNγ on the epithelial barrier is significantly affected by other immune mediators. Its pro-inflammatory effects increase in the presence of TNF-α, whereas SOCS1 inhibits its signaling pathways (Larkin, 2013). IFNγ disrupts barrier integrity by downregulating junction proteins and promoting bacterial translocation (Clark, 2005; Han, 2019). IFNγ stimulation induces CXCL9, CXCL10, and CXCL11 secretion from epithelial cells, attracting cytotoxic T cells and NK cells that exacerbate inflammation (Suzuki, 2007; Dwinell, 2001; Kulkarni, 2017).

TNFa is the most targeted factor for therapeutics in IBD treatment, with the majority of IBD treatments being biologics against TNFa; as many as a third of IBD patients are primary non-responders to TNFa and about half of those that initially respond to treatment end up secondary non-responders (Kumar, 2024). Despite identifying TNFa as a prominent contributor to IBD pathology, targeting TNFa alone is not an adequate treatment. Targeting CXCR3 and its ligand CXCL10 is a promising approach in chemokine therapy for IBD (Trivedi, 2018). High CXCR3 expression on activated T-cells has been observed in IBD patient tissue biopsies, with increased CXCL10 potentially leading to epithelial cell death. The chemokines CXCL9, CXCL10, and CXCL11 are all elevated in preclinical models of UC, with CXCR3 ligands attracting CD4+CXCR3+ T cells to the epithelial barrier, exacerbating inflammation (Suzuki, 2007; Dwinell, 2001; Kulkarni, 2017). While these chemokines are often grouped, they play different roles in the immune response. CXCL10 only binds to CXCR3 and is noted as particularly important in its involvement in autoimmunity (Christen, 2003).

While there are several biologics available for the treatment of IBD, there remains a significant unmet need in the therapeutic landscape. These biologics, though effective for many patients, do not offer relief for all. Many patients either do not respond to current treatments or lose responsiveness over time. This highlights the need for the development of novel therapeutics that can offer broader efficacy and improved safety profiles, targeting the diverse pathways involved in IBD pathogenesis.

## Materials and Methods

### Animal Procurement and Housing

Male and female B6 mice aged 8-12 weeks were obtained from Jackson Laboratories. Animals were housed under controlled conditions with a 12-hour light/dark cycle, temperature maintained at 22 ± 2°C. They were given access to standard laboratory chow and water ad libitum throughout the study period.

All animal experiments were conducted in strict accordance with the recommendations in the Guide for the Care and Use of Laboratory Animals of the National Institutes of Health. The protocol was approved by the Institutional Animal Care and Use Committee (IACUC) of the host institution. All efforts were made to minimize animal suffering and to reduce the number of animals used.

### Crypt Isolation and Organoid Culture Maintenance

Intestinal crypt isolation was performed using B6 mice. The intestine was removed and flushed with Dulbecco’s Phosphate Buffered Saline (GIBCO) containing 0.1% Bovine Serum Albumin (DPBS 0.1% BSA (Sigma Aldrich)). The intestines were then sectioned into 3 mm pieces and washed with DPBS 0.1% BSA. The tissue segments were incubated at 37°C with Gentle Cell Dissociation Reagent (Stemcell Technologies) for 5 minutes with shaking at 100 rpm. Subsequently, the tissue was subjected to vigorous manual shaking for 15 seconds and then briefly vortexed. The supernatant was discarded, and fresh 37°C Gentle Cell Dissociation Reagent was added before placing the tissue back in the shaker for an additional 15 minutes. This was followed by a 15-20 minute incubation at room temperature without shaking. The tissue was then vigorously shaken by hand for 15 seconds, and the supernatant was filtered through a 100 µm cell strainer. DPBS 0.1% BSA was added, and the process was repeated twice more. The final supernatant, containing the isolated crypts, was passed through a 70 µm filter. The crypts were resuspended in a solution of Matrigel (Corning Cat #354248) and Murine IntestiCult (Stemcell Technologies) at a concentration of 60 crypts per 10 µl and seeded into a 96-well plate. Matrigel was diluted to 1x concentration using DMEM (Corning) supplemented with 50ug/mL gentamycin (Gibco).

Organoid cultures were maintained by refreshing the media every two to three days. The cultures were initially passaged on day six following seeding, with subsequent passages taking place every three to four days thereafter. Experiments were conducted on organoids within three passages from primary isolation to ensure consistency and viability of the samples.

### Mode-k Cell Culture

Mode-K cells were retrieved from liquid nitrogen storage and cultured in T75 flasks. The culture media were replenished every 2 to 3 days, and the cells were passaged at 6-day intervals. The Mode-K culture medium consisted of Dulbecco’s Modified Eagle Medium (DMEM) supplemented with 5% fetal bovine serum (FBS), 1% HEPES buffer (Gibco), 1% sodium pyruvate (Corning) 0.1% beta-mercaptoethanol (Gibco), and penicillin/streptomycin. For experiments, Mode-K cells were seeded into 96 or 12 well TC treated plates.

### Propidium Iodide/ Hoechst Live Dead Staining

Organoids and Mode-K cells were incubated at 37°C with 5% CO₂ in culture media supplemented with 5 µg/mL PureBlu Hoechst (Bio-Rad) and 5 µg/mL Propidium Iodide, for a duration of 45 minutes. Following incubation, the culture media was discarded, and the cells were washed with pre-warmed PBS. The plates were then analyzed using a Cytation 5 imaging reader (Agilent Technologies). A 7×7 area scan was conducted for fluorescence detection at the following settings: Excitation at 535/20 nm and Emission at 617/20 nm for Propidium Iodide, and Excitation at 361/20 nm and Emission at 486/20 nm for Hoechst staining.

### ELISA

ELISAs for CXCL10 and TNF-alpha were performed using Duo-set kits from R&D Systems following the manufacturer’s guidelines. The assays included multiple wash steps performed on a designated machine with a bottom wash followed by three additional washes. The blocking step involved reagent diluents at 5% concentration. Absorbance was measured using a Cytation 5 imaging reader (Agilent Technologies) at 450 nm, with 570 nm serving as the reference wavelength.

### Immunofluorescence Microscopy

The culture medium was discarded, and the cells were washed with PBS. Cells were then fixed and permeabilized in 100% ice-cold methanol at −20°C for 4-5 minutes. After the removal of methanol, cells were washed again with PBS and subsequently blocked with 5% goat serum for 1 hour at room temperature with gentle shaking. Following another PBS wash, primary antibodies (Phospho-Stat3 (Tyr705) (D3A7) XP® Rabbit mAb #9145 and Phospho-Stat1 (Tyr701) (58D6) Rabbit mAb #9167, Cell Signaling) were diluted at a ratio of 1:200 and applied to the cells, which were then left to incubate overnight at 4°C.

The next day, the primary antibody solution was removed, and cells were washed with PBS. Secondary antibodies (Goat anti-Rabbit IgG (H+L) Highly Cross-Adsorbed Secondary Antibody, Alexa Fluor™ Plus 555, Invitrogen) were diluted between 1:500 and 1:1000 and added to the wells. Cells were incubated for 1 hour with gentle shaking in the dark. Afterward, the secondary antibody was discarded, and cells were stained with DAPI in PBS for 15 minutes with shaking at room temperature. A final wash with PBS was performed before acquiring images on a Cytation 5 imaging system. Analysis was performed using Gen5 Software by Biotek (Agilent).

### RNA Isolation and cDNA Synthesis

Total RNA was extracted from tissue samples using Qiagen’s RNeasy Mini Spin Columns, incorporating an on-column DNase digestion to eliminate genomic DNA contamination. cDNA was synthesized using the SuperScript IV Reverse Transcriptase kit from Thermo Fisher Scientific, with a modified protocol that included an extended incubation at 50°C to improve transcript yield.

### QPCR

Gene expression levels were quantified by qPCR using iTaq Universal SYBR Green Supermix (Bio-Rad). Specific primers for each gene, detailed in Table S1, were used. Reactions were performed on a QuantStudio 5 Real-Time PCR System according to the standard SYBR Green protocol. All samples were analyzed in duplicate to ensure accuracy and reproducibility.

### Scratch Assay

The scratch assay was performed on mode-k cells that were cultured to near or full confluency in 96-well plates. A sterile 1000 µl pipette tip connected to a vacuum system was used to create a uniform scratch across the cell monolayer. Subsequent to the scratch, cells were maintained in their growth medium under standard culture conditions. Brightfield microscopy images were captured at 0 and 36 hours post-scratch using a Cytation 5 Imaging Reader. Image analysis for quantifying wound closure was conducted using ImageJ software, employing the Scratch Wound Assay Macro developed by the MRI Group. This analysis provided quantitative data on cell migration and wound healing over time.

### Statistical Analysis

All statistical analysis were performed using Graphpad Prism 10.1.2. Ordinary One-Way ANOVA followed by Tukey’s Multi-Comparisons tests. T-test were used for groups of two. ns=not significant

### Use of AI Language Model

ChatGPT, an AI language model developed by OpenAI, was utilized during the preparation of this draft. The general workflow was as follows: First, a human writes a draft with all the information. The draft is given to ChatGPT with the prompt: “Please rewrite the following so it flows better. Put the in-text citations into APA format. Do not remove or add any information. In-text citations need to remain with their facts. Please do not use the word “pivotal.” Please correct spelling and grammar mistakes. Avoid introductory clauses”. The author would take the rewritten content and edit what ChatGPT wrote because, invariably, it has failed some of the requests in the prompt. In a second method used, ChatGPT would be given the main ideas wanted in a section and asked to write that section. The output was typically exceptionally unfit, but the general structure was useful. So, the author would use the structure to write their content and then employ method one. The author does not think the process was faster than writing without ai assistance, but the manuscript is probably less painful for others to read.

## Results

REG3b, REG3g, TNFa, and CXCL10, selected for their relevance to inflammatory bowel disease (IBD) and robust expression in small intestine (SI) derived organoids, were investigated in response to stimulation by IFNγ and IL-22 (Fig.S1).

### Modulation of inflammatory Profile: Downregulation of Antimicrobial Peptides and Upregulation of Pro-inflammatory Markers CXCL10 and TNFα

To explore the co-stimulatory effects of IFNγ and IL-22 on Intestinal epithelial cells, we stimulated small intestine-derived organoids with cytokines for 24 hours and evaluated gene expression by QPCR. After 48 hours of stimulation, we evaluated protein levels in the supernatant by ELISA. IL-22 stimulation increased REG3b and REG3y expression at doses of 1ng/mL and 10ng/mL (Fig1.A-B). When 1ng/mL of IFNγ was added to the organoids along with IL-22, a significant reduction in REG3b and REG3y expression was observed (Fig.1A-B).

**Figure 1.**
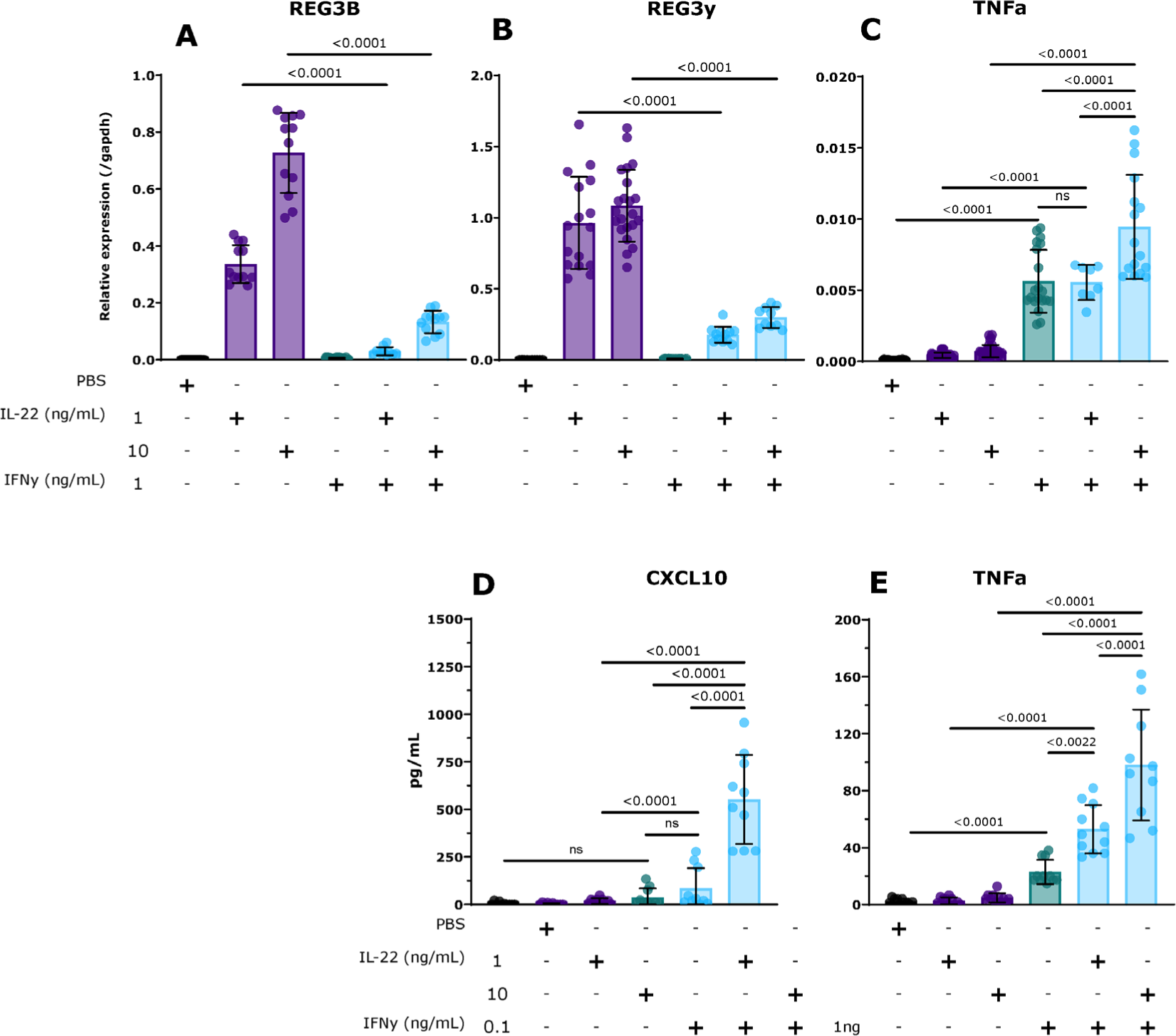
IFNy and IL-22 Co-stimulation Effects on SI Organoids Gene Expression and Protein Levels. Organoids were stimulated for 24 hour and evaluated for gene expression by QPCR. Protein expression was assessed at 48 hours via ELISA. (A,B) IFNy dampens IL-22 stimulated upregulation of AMPs. (C,E) IL-22 amplifies IFNy-induced TNFa gene expression and protein concentration in supernatant (D) IL-22 and IFNy synergize increasing CXCL10 protein concentration in a dose-dependent manner. QPCR samples ran in technical duplicates and collected from >3 experiments. ELISA data point represent independent biological samples collected from ≥ 3 experiments. Ordinary one-way ANOVA followed by Tukey’s multiple comparisons test and expressed as mean +/− SD.

Organoids co-stimulated with 1ng/mL of IFNγ and 1ng/mL of IL-22 did not express TNFα significantly more than organoids stimulated with IFNγ alone (Fig.1C). In contrast, combining 1ng/mL IFNγ with 10ng/mL IL-22 significantly increased TNFα expression (Fig.1C). Next we confirmed the QPCR data by ELISA. When 1ng/mL IL-22 was added to the 1ng/mL IFNγ stimulation, TNFα levels showed a significant increase compared to the levels induced by either cytokine alone (Fig.1E). Further enhancement of TNFα expression was observed when the concentration of IL-22 was increased to 10ng/mL while maintaining IFNγ at 1ng/mL. The TNFα levels in this condition were significantly higher than those observed with the lower dose combination of 1ng/mL IL-22 and 1ng/mL IFNγ (Fig.1E).

Previous studies have shown IFNy and TNFa co-stimulation of epithelial cells upregulates ISGs including CXCL10 (Oslund, 2014). Having confirmed a significant increase of TNFa in the supernatant of the organoids co-stimulated with IFNy and IL-22, we predicted these conditions would increase concentrations of CXCL10. To test this we employed the same experimental design as when evaluating for TNFa.

Administration of 0.1ng/mL IFNγ alone did not induce a significant amount of CXCL10 compared to PBS control (Fig.1D). When 1ng/mL IL-22 was combined with 0.1ng/mL IFNγ, there was a significant enhancement in CXCL10 levels compared to the stimulation with IL-22 alone (Fig.1D). However, this combination did not result in a statistically significant difference from the CXCL10 levels induced by IFNγ alone. The most notable increase in CXCL10 protein levels occurred when 0.1ng/mL IFNγ was combined with 10ng/mL IL-22 (Fig.1D). This combination significantly elevated CXCL10 induction above all other tested conditions.

### Increased Cell Death in Response to IFNγ and IL-22 Co-Stimulation Partially Due to TNFa

An essential aspect of epithelial barrier defense and IBD pathology is cell death. A breakdown in the barrier allows for the contents of the lumen and the immune system to come into contact, risking elevated immune activation and perpetuation of inflammation (Odenwald, 2017). IFNγ is known to induce cell death through apoptosis/necroptosis. TNFa has also been studied as a factor capable of inducing cell death in epithelial cells (Woznicki, 2021). Given the results showing an increase in TNFa in the combined co-stimulatory conditions, we were interested in exploring possible effects on cell viability. We employed a propidium iodide and Hoechst staining assay (modified from Bode 2019) to quantitatively measure cell death in mouse intestinal organoids following exposure to varying concentrations of the cytokines over 48 hours.

The percentage of cell death in the organoids demonstrated a dose-dependent increase when stimulated with IFNγ. The cell death rate escalated with higher doses of IFNγ and reached a plateau at 10ng/mL, where it stabilized at approximately 60% mortality (Fig.2B). In contrast, organoids stimulated with IL-22 exhibited a different pattern of cell death. The mortality rate increased with escalating doses of IL-22 but reached a plateau at a lower concentration of 5ng/mL, where cell death accounted for about 20% of the population (Fig.2C).

**Figure 2.**
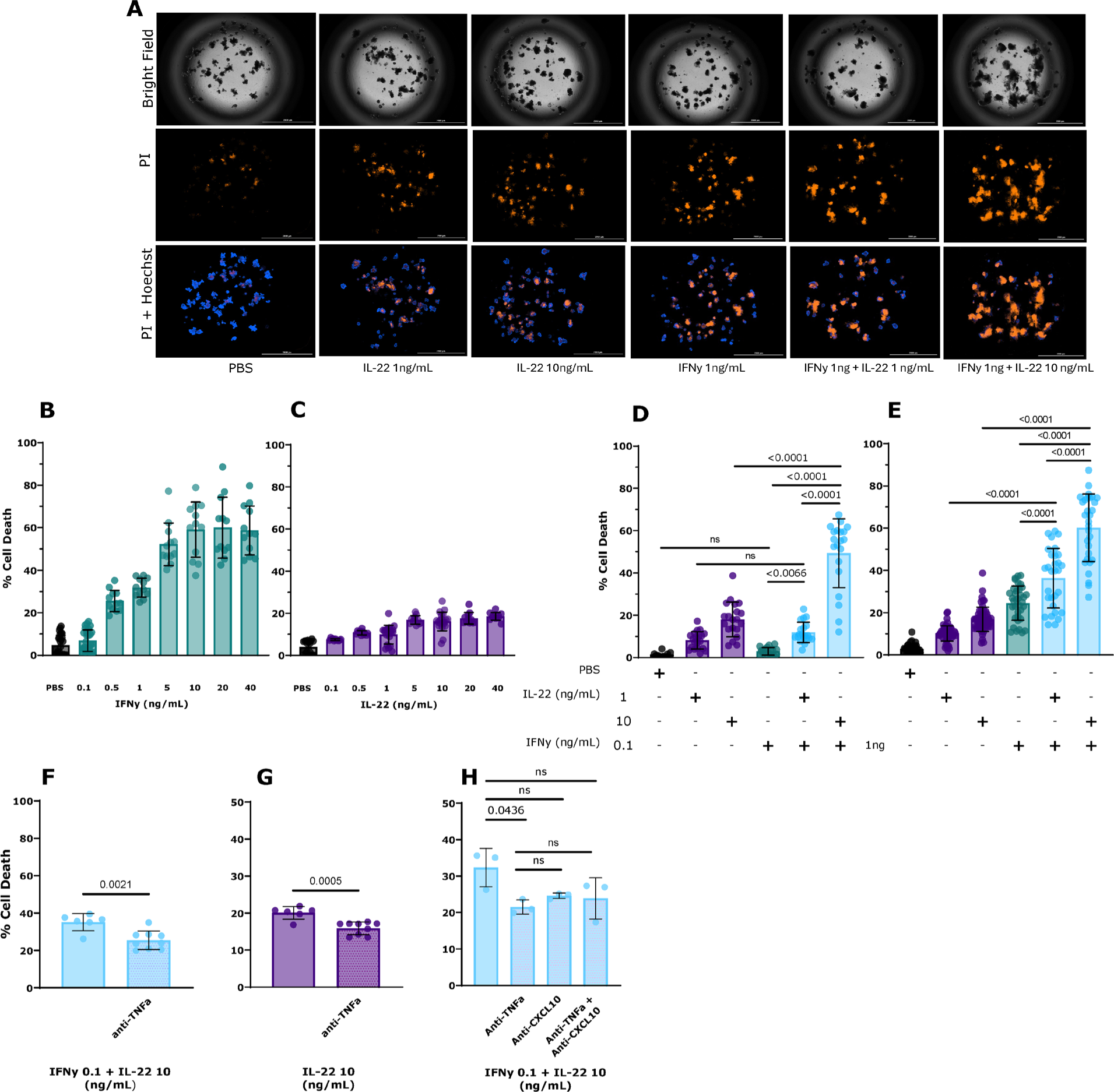
Cytokine Stimulation on Organoid Cell Viability. SI organoids were stimulated for 48 hours then evaluated by PI/H for cell death*. (*A) PI/H images of cytokine stimulated Organoids (B,C) Increasing concentration of cytokine increases cell death. (D,E) IL-22 and IFNy synergize, enhancing cell death in organoids. (F-H) Anti-TNFa partially ameliorates cytokine-induced cell death. Data points are representative of one well and were collected over ≥ 3 experiments (H’s data is from 1 experiment). Significance was determined using Ordinary one-way ANOVA followed by Tukey’s multiple comparisons test and expressed as mean +/− SD. Groups of two evaluated by T-tests.

Organoids treated with 0.1ng/mL IFNγ alone caused minimal cell death (Fig.2D). Co-stimulation with IL-22 at 1ng/mL and IFNγ at 0.1ng/mL resulted in a significant increase in cell death compared to IFNγ alone (Fig.2D). A more pronounced cytotoxic effect was observed when the concentration of IL-22 was increased to 10ng/mL while maintaining IFNγ at 0.1ng/mL. Under these conditions, approximately 50% of the organoid(s) underwent cell death after 48 hours of stimulation (Fig.2D).

The same experiment was conducted but with IFNγ at 1ng/mL, with similar results. IFNγ alone induced substantial cell death, which was augmented by co-stimulation with IL-22 (Fig.2E). In this experiment, the co-stimulatory conditions exhibited significantly more cell death than the cytokines individually (Fig.2E). Additionally, the higher dose of IL-22 resulted in more cell death than the lower dose of IL-22 in the combined conditions (Fig.2E).

Next, we investigated TNFa as a possible contributor to the increase in cell death observed in the co-stimulatory conditions. Organoids were stimulated with 0.1ng/mL of IFNγ and 10ng/mL of IL-22 for 48 hours. One group of organoids was also treated with anti-TNFα. The anti-TNFα-treated group exhibited significantly reduced cell death compared to the group that did not receive anti-TNFα (Fig.2F).

In the IL-22 experiments analyzing cell death, at varying doses, after a concentration of 5ng/ml, around 20%, cell death invariably occurs in organoids (Fig.2C). We predicted this cytotoxicity could be partially attributed to TNFa, so we mirrored the last experiment. However, we focused only on IL-22 10ng/mL stimulated organoids. The group treated with anti-TNFα showed significantly less cell death than the untreated group (Fig.2G).

Since we also observed an increase in CXCL10 and previous studies have shown that CXCL10 can induce cell death (Singh, 2009), we explored CXCL10’s potential contribution to cell death. We tested responses to combined IL-22 and IFNγ co-stimulation under four different conditions: no inhibitors, anti-TNFα, anti-CXCL10, and a combination of both inhibitors. The introduction of anti-TNFα alone markedly decreased cell death compared to the untreated control (Fig.2H). Conversely, anti-CXCL10, whether alone or combined with anti-TNFα, did not significantly affect cell death (Fig.2H).

### IL-22 and IFNγ Co-stimulation Slows Wound Healing

Further investigating the effects of co-stimulation of IFNγ and IL22 and barrier integrity, we used mode-k cells to perform a scratch/wound healing assay. Previous studies have found that IL-22 generally accelerates the rate of wound closure through proliferation and migration (Avitibile, 2015). IFNγ in wound healing assays promotes closure through migration in certain cell types at specific doses (He, 2017).

In this experiment, mode-k cells were grown to/near confluence, a scratch was made in the monolayer, and after 36 hours, the scratch was assessed for percent closure. Prior to the scratch, the cells were pre-stimulated with cytokines.

IL-22 at higher doses (10ng/mL) accelerated wound closure compared to PBS (Fig.3C), while IFNγ slowed wound closure (Fig.3D). Co-stimulation with IL-22 and IFNγ impaired wound healing more than IFNγ alone (Fig.3D).

**Figure 3.**
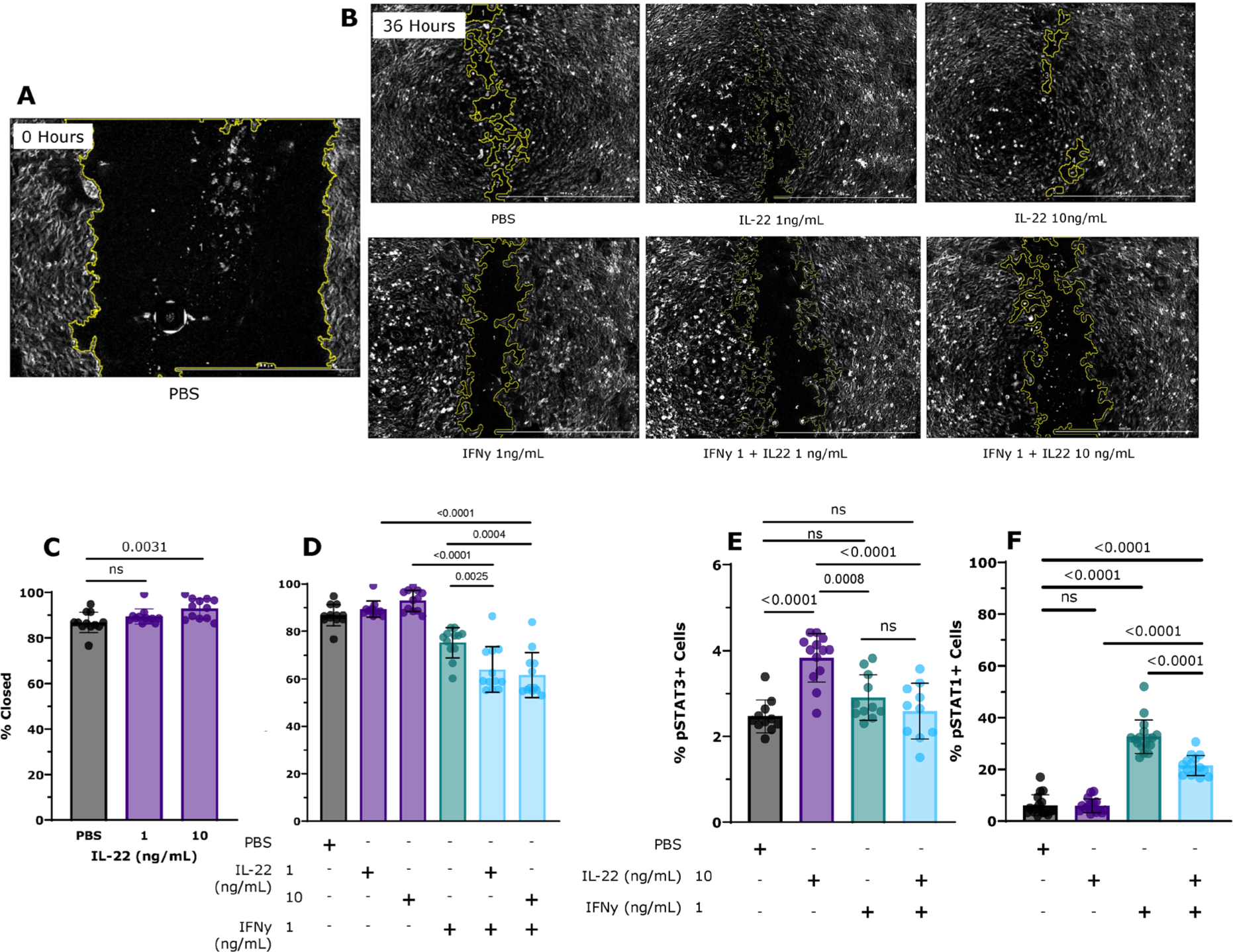
IL-22 and IFNy Co-stimulation in Wound Healing. Scratch assay was performed on mode-k cells with 24 hours of stimulation prior to (and during) the scratch and evaluation 36 hours post scratch. (A). Digital phase contrast image of PBS scratch at 0 hours. (B.) Digital Phase contrast images of all conditions at 36 hours post scratch. (C). Higher dose IL-22 accelerates wound closure compared to PBS. (D) IL-22 and IFNy co-stimulation slows wound closure. Mode-k cells’ pSTAT3 and pSTAT1 was evaluated by immunofluorescence after 24 hours of stimulation (E). IL-22 increased pSTAT3. Co-stimulation of IL-22 with IFNy significantly reduced pSTAT3 levels, compared to Il-22 alone. (F) IFNy stimulation significantly increases pSTAT1. Co-stimulation with IL-22 significantly decreases pSTAT1, compared to IFNy alone. C,D data points represent 1 well collected from 3 experiments. E,F data points represent 1 image collected from 2 experiments. Significance was determined using Ordinary one-way ANOVA followed by Tukey’s multiple comparisons test and expressed as mean +/− SD.

Pickert 2009 linked IL-22 and phosphorylated STAT3 (pSTAT3) to mucosal wound healing, with the absence of STAT3 in IECs significantly impairing healing capabilities. Given the results of the scratch assays, we predicted IL-22 would enhance phosphorylated STAT3, and co-stimulation with IFNγ would negate IL-22-stimulated elevated pSTAT3 levels. Mode-K cells were stimulated with cytokines for 24 hours and then evaluated for pSTAT3 and pSTAT1 via immunofluorescence.

When mode-K cells were stimulated with IL-22 at 10ng/mL, the number of pSTAT3-positive cells compared to the baseline and other experimental groups was pronouncedly increased (Fig.3E) IF Images (Fig.S1A). Co-stimulation of cells with IL-22 at 10ng/mL and IFNγ at 1ng/mL reduced the number of pSTAT3-positive cells, reverting to baseline levels (Fig.3E).

Stimulation of mode-K cells with IFNγ alone significantly increased the presence of pSTAT1-positive cells (Fig.3F). IL-22 at 10ng/mL was introduced alongside IFNγ, there was a noticeable reduction in pSTAT1-positive cells, although levels remained above baseline (Fig.3F) IF Images (Fig.S1B).

## Discussion

This study investigated the interactions between IL-22 and IFNγ on epithelial cells, particularly concerning cytokine-driven gene expression and cellular responses, including cytotoxicity and wound healing. The findings reveal intricate interactions between these cytokines that affect cellular processes relevant to IBD pathogenesis.

IL-22 and IFNγ traditionally have opposing roles, with IL-22 typically promoting regenerative responses and IFNγ mediating inflammatory actions. This study suggests that co-stimulation with IFNγ can convert the usually protective effects of IL-22 into a pro-inflammatory response.

IFNγ dampens IL-22-induced REG3b and REG3y expression, which is crucial for mucosal healing and antimicrobial defense. This interaction suggests a dominant inhibitory role of IFNγ over the beneficial effects of IL-22 in epithelial repair and maintenance. The synergistic effect of IL-22 and IFNγ on the induction of TNFα and CXCL10 further demonstrates the complexity of cytokine interactions. Surprisingly, higher doses of IL-22 further enhanced the induction of TNFa and CXCL10. The elevated levels of TNFa are notably problematic as elevated TNFa is one of the most prominent characteristics of IBD. TNFα synergizes with IFNγ, acting on epithelial cells which produce elevated levels of CXCL9, CXCL10, CXCL11, and TNFα. CXCL9, CXCL10, and CXCL11, through chemotaxis, attract CD8 cytotoxic T cells and NK cells, which produce IFNγ and TNFα, creating a feedforward loop that further promotes a pro-inflammatory environment (Suzuki, 2007; Dwinell; 2001 Kulkarni 2017). CXCR3 and its ligand, particularly CXCL10, are being investigated as therapeutic targets in autoimmunity (Christen, 2003). The cytotoxic effects induced by co-stimulation with IFNy and IL-22 in intestinal organoids, and the partial mitigation of these effects by anti-TNFα treatment, underscore TNFα’s critical role in cytokine-induced epithelial damage. This finding is consistent with the known pro-apoptotic functions of TNFα and supports ongoing therapeutic strategies targeting TNFα in IBD. The lack of significant impact of anti-CXCL10 treatment on cell viability indicates a secondary role of CXCL10 in cytokine-induced cytotoxicity despite its upregulation in inflammatory conditions.

In the context of epithelial barrier integrity, the inhibition of wound closure by IFNγ highlights the challenges in managing IBD, where healing of mucosal lesions is crucial. IFNy and IL-22 co-stimulation further inhibiting wound closure is particularly problematic, as IL-22 is critical in wound healing.

Lastly, the alterations in STAT signaling induced by cytokine co-stimulation provide a molecular basis for the observed cellular responses. The reduction of pSTAT3-positive cells in the presence of IFNγ, even with high levels of IL-22, reveals a competitive antagonism between these cytokines at the signaling level. Conversely, the decrease in pSTAT1 expression with IL-22 addition suggests a partial counter-regulatory mechanism, though insufficient to reverse IFNγ effects fully.

## Conclusion

An essential insight from this study, with significant therapeutic implications, is the critical role of cytokine dosage. We observed that problematic interactions between these cytokines predominantly occurred at higher concentrations and in combination. This underscores the importance of not only discerning the interactions between cytokines but also understanding how these interactions vary with concentration. Such knowledge can refine our approaches to treatment. This nuanced understanding of dosage effects highlights the need for precision in cytokine modulation strategies in clinical and research settings.

The unexpected heightened pro-inflammatory response to co-stimulation with IL-22 and IFNγ in this study emphasizes the need for therapeutic strategies that consider the context-dependent effects of cytokines in IBD. Modulating these cytokines to avoid unintended pro-inflammatory responses could enhance therapeutic efficacy and minimize adverse effects. Future therapies might benefit from targeting specific cytokine interactions to better control inflammation without disrupting the essential protective roles of cytokines like IL-22 in the gut epithelium.

## References

1. Abraham, C., & Cho, J. H. (2009). Inflammatory bowel disease. *The New England Journal of Medicine, 361*(21), 20662078. 10.1056/NEJMra0804647

2. Aparicio-Domingo, P., Romera-Hernández, M., Karrich, J. J., Cornelissen, F., Papazian, N., Lindenbergh-Kortleve, D. J., Butler, J. A., Boon, L., Coles, M. C., Samsom, J. N., & Cupedo, T. (2015). Type 3 innate lymphoid cells maintain intestinal epithelial stem cells after tissue damage. *Journal of Experimental Medicine, 212*(11), 1783–1791. 10.1084/jem.20150318

3. Christen, U., McGavern, D. B., Luster, A. D., von Herrath, M. G., & Oldstone, M. B. (2003). Among CXCR3 chemokines, IFN-gamma-inducible protein of 10 kDa (CXC chemokine ligand (CXCL)10) but not monokine induced by IFN-gamma (CXCL9) imprints a pattern for the subsequent development of autoimmune disease. *Journal of Immunology, 171*, 6838–6845.

4. Chiba, M., Nakane, K., & Komatsu, M. (2019). Westernized diet is the most ubiquitous environmental factor in inflammatory bowel disease. *The Permanente Journal, 23*, 18–107. 10.7812/TPP/18-107

5. Clark, E., Hoare, C., Tanianis-Hughes, J., Carlson, G. L., & Warhurst, G. (2005). Interferon gamma induces translocation of commensal Escherichia coli across gut epithelial cells via a lipid raft-mediated process. *Gastroenterology, 128*(5), 1258–1267. 10.1053/j.gastro.2005.01.046

6. Conlon, T. M., Knolle, P. A., & Yildirim, A. (2021). Local tissue development of type 1 innate lymphoid cells: guided by interferon-gamma. *Signal Transduction and Targeted Therapy, 6*, 287. 10.1038/s41392-021-00705-1

7. Dwinell, M. B., Lügering, N., Eckmann, L., & Kagnoff, M. F. (2001). Regulated production of interferon-inducible T-cell chemoattractants by human intestinal epithelial cells. *Gastroenterology, 120*, 49–59.

8. Fish, S. M., Proujansky, R., & Reenstra, W. W. (1999). Synergistic effects of interferon gamma and tumour necrosis factor alpha on T84 cell function. *Gut, 45*(2), 191–198. 10.1136/gut.45.2.191

9. Gao, S., Li, Y., Wu, D., Jiao, N., Yang, L., Zhao, R., Xu, Z., Chen, W., Lin, X., Cheng, S., Zhu, L., Lan, P., & Zhu, R. (2022). IBD subtype-regulators IFNG and GBP5 identified by causal inference drive more intense innate immunity and inflammatory responses in CD than those in UC. *Frontiers in Pharmacology, 13*, 869200. 10.3389/fphar.2022.869200

10. Gonsky, R., Deem, R. L., Landers, C. J., Haritunians, T., Yang, S., & Targan, S. R. (2014). IFNG rs1861494 polymorphism is associated with IBD disease severity and functional changes in both IFNG methylation and protein secretion. *Inflammatory Bowel Diseases, 20*(10), 1794–801. 10.1097/MIB.0000000000000172

10. Hu, Y., Chen, Z., Xu, C., Kan, S., & Chen, D. (2022). Disturbances of the Gut Microbiota and Microbiota-Derived Metabolites in Inflammatory Bowel Disease. *Nutrients, 14*(23), 5140. 10.3390/nu14235140

11. Keir, M., Yi, Y., Lu, T., & Ghilardi, N. (2020). The role of IL-22 in intestinal health and disease. *Journal of Experimental Medicine, 217*(3), e20192195. 10.1084/jem.20192195

12. Kim, S., Nowakowska, A., Kim, Y. B., & Shin, H. Y. (2022). Integrated CRISPR-Cas9 system-mediated knockout of IFN-γ and IFN-γ receptor 1 in the Vero cell line promotes viral susceptibility. *International Journal of Molecular Sciences, 23*(15), 8217. 10.3390/ijms23158217

13. Kulkarni, N., Manisha, P., & Girdhari, L. (2017). Role of chemokine receptors and intestinal epithelial cells in the mucosal inflammation and tolerance. *Journal of Leukocyte Biology, 101*, 377–394.

14. Kumar, M., Murugesan, S., Ibrahim, N., et al. (2024). Predictive biomarkers for anti-TNF alpha therapy in IBD patients. *Journal of Translational Medicine, 22*, 284. 10.1186/s12967-024-05058-1

15. McDowell, C., Farooq, U., & Haseeb, M. (2023). Inflammatory Bowel Disease. In StatPearls [Internet]. Treasure Island (FL): StatPearls Publishing. Available from: https://www.ncbi.nlm.nih.gov/books/NBK470312/

16. Mizoguchi, A., Yano, A., Himuro, H., et al. (2018). Clinical importance of IL-22 cascade in IBD. *Journal of Gastroenterology, 53*, 465–474. 10.1007/s00535-017-1401-7

17. Murray, P. D., McGavern, D. B., Pease, L. R., & Rodriguez, M. (2002). Cellular sources and targets of IFN-gamma-mediated protection against viral demyelination and neurological deficits. *European Journal of Immunology, 32*(3), 606–615. 10.1002/1521-4141(200203)32:3<606::AID-IMMU606>3.0.CO;2-D

18. Ng, S. C., Shi, H. Y., Hamidi, N., Underwood, F. E., Tang, W., Benchimol, E. I., Panaccione, R., Ghosh, S., Wu, J. C. Y., Chan, F. K. L., Sung, J. J. Y., & Kaplan, G. G. (2017). Worldwide incidence and prevalence of inflammatory bowel disease in the 21st century: A systematic review of population-based studies. *The Lancet, 390*(10114), 2769–2778. 10.1016/S0140-6736(17)32448-0

19. Oslund KL, Zhou X, Lee B, Zhu L, Duong T, Shih R, Baumgarth N, Hung LY, Wu R, Chen Y. Synergistic up-regulation of CXCL10 by virus and IFN γ in human airway epithelial cells. PLoS One. 2014 Jul 17;9(7):e100978. doi: 10.1371/journal.pone.0100978. PMID: 25033426; PMCID: PMC4102466.

20. OpenAI ChatGPT. (2023). AI language model developed by OpenAI.

21. Perez-Toledo, M., Beristain-Covarrubias, N., Channell, W. M., Hitchcock, J. R., Cook, C. N., Coughlan, R. E., Bobat, S., Jones, N. D., Nakamura, K., Ross, E. A., Rossiter, A. E., Rooke, J., Garcia-Gimenez, A., Jossi, S., Persaud, R. R., Marcial-Juarez, E., Flores-Langarica, A., Henderson, I. R., Withers, D. R., Watson, S. P., & Cunningham, A. F. (2020). Mice deficient in T-bet form inducible NO synthase-positive granulomas that fail to constrain Salmonella. *The Journal of Immunology, 205*(3), 708–719. 10.4049/jimmunol.2000089

22. Platanias, L. (2005). Mechanisms of type-I- and type-II-interferon-mediated signalling. *Nature Reviews Immunology, 5*, 375–386. 10.1038/nri1604

23. Suzuki, K., Kawauchi, Y., Palaniyandi, S. S., Veeraveedu, P. T., Fujii, M., Yamagiwa, S., Yoneyama, H., Han, G. D., Kawachi, H., Okada, Y., Ajioka, Y., Watanabe, K., Hosono, M., Asakura, H., Aoyagi, Y., & Narumi, S. (2007). Blockade of interferon-gamma-inducible protein-10 attenuates chronic experimental colitis by blocking cellular trafficking and protecting intestinal epithelial cells. *Pathology International, 57*, 413–420.

24. Trivedi, P. J., & Adams, D. H. (2018). Chemokines and chemokine receptors as therapeutic targets in inflammatory bowel disease; pitfalls and promise. *Journal of Crohn’s and Colitis, 12*(suppl_2), S641–S652. 10.1093/ecco-jcc/jjx145

25. Zheng, Y., Valdez, P. A., Danilenko, D. M., Hu, Y., Sa, S. M., Gong, Q., Abbas, A. R., Modrusan, Z., Ghilardi, N., de Sauvage, F. J., & Ouyang, W. (2008). Interleukin-22 mediates early host defense against attaching and effacing bacterial pathogens. *Nature Medicine, 14*(3), 282–289. 10.1038/nm1720

