## Supplemental Data for "Modulatory Effects of IFN-γ and IL-22 on Inflammatory Signaling and Cellular Responses in Intestinal Epithelial Cells"

Table S1

| Primers | 5’ Forward 3’ | 5’Reverse3’ |
|---|---|---|
| **GAPDH** | CGACTTCAACAGCAACTCCCACTCTTCC | TGGGTGGTCCAGGGTTTCTTACTCTT |
| **REG3y** | TTCCTGTCCTCCATGATCAAAA | CATCCACCTCTGTTGGGTTCA |
| **REG3b** | ATGCTGCTCTCCTGCCTGATG | CTAATGCGTGCGGAGGGTATATTC |
| **CXCL10** | GGATGGCTGTCCTAGCTCTG | TGAGCTAGGGAGGACAAGGA |
| **TNFa** | CGATCACCCCGAAGTTCAGTA | CAGGCGGTGCCTATGTCTC |

**Figure S1**

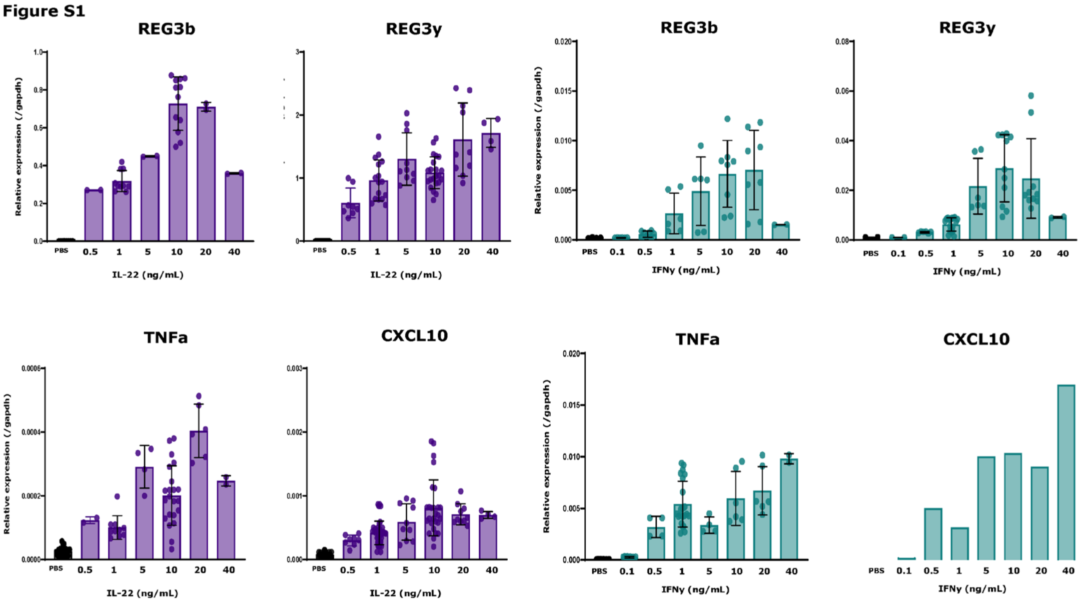


***Figure S1. SI Organoids gene expression in response to IFNy and IL-22 stimulation***

**Figure S2**

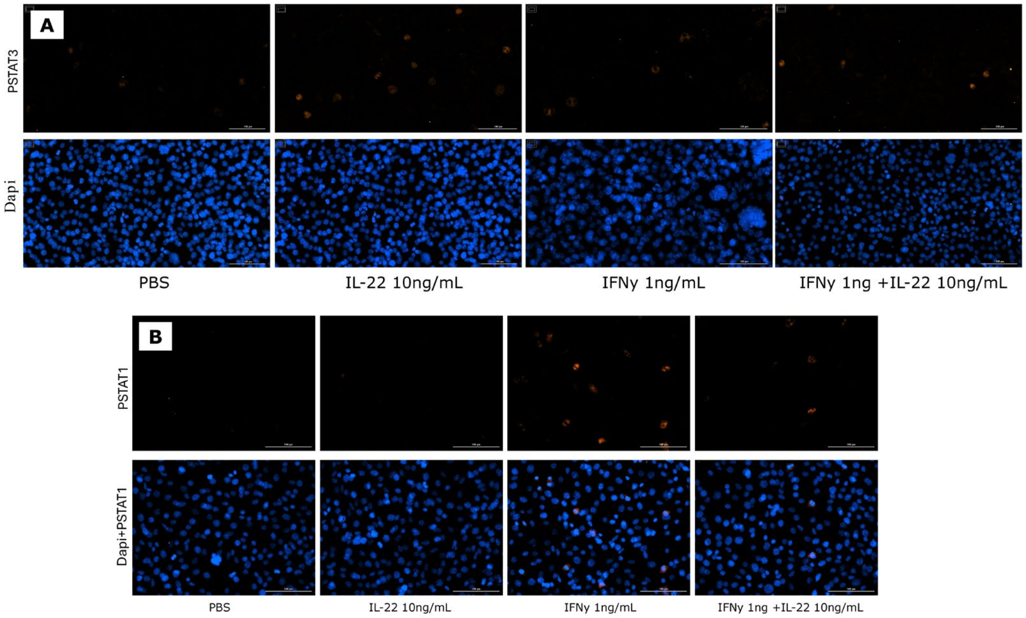


***Figure S2. Phosphorylated STAT Immunofluorescence.*** (A) pSTAT3 over all conditions. (B) pSTAT1 over all conditions
